# Environmental Enrichment Attenuates Fentanyl-Seeking Behavior and Protects Against Stress-Induced Reinstatement in Both Male and Female Rats

**DOI:** 10.64898/2025.12.02.691896

**Authors:** Jessica A. Higginbotham, Joanna J. Dearman, Hasan Maqbool, Mateo Pujol, Jose A. Moron

## Abstract

Environmental enrichment (EE) reduces vulnerability to multiple drugs of abuse, yet its impact on fentanyl use and relapse-like behavior remains unclear. Here, we tested whether long-term, non-social, object-based EE alters fentanyl self-administration, extinction, and stress-induced reinstatement in male and female rats. Rats were individually housed in either standard non-enriched (NE) conditions or in EE cages containing a rotating set of novel objects beginning at least three days prior to self-administration. EE did not impact acquisition of fentanyl self-administration but reduced fentanyl intake during maintenance of self-administration and reduced the persistence of drug-seeking in extinction. Following extinction, yohimbine robustly reinstated drug-seeking behavior in NE rats but reinstatement in EE rats was markedly attenuated, indicating reduced sensitivity to stress-induced relapse triggers. Circulating corticosterone levels were lower in EE rats across the experiment and were positively correlated with reinstatement responding, suggesting that enrichment’s protective effects may be mediated in part by reduced hypothalamic-pituitary-adrenal (HPA) axis activation. These findings demonstrate that object-based EE, even in the absence of social enrichment, is sufficient to blunt fentanyl use, facilitate extinction, and constrain stress-induced reinstatement. The results highlight the role of environmental context and stress regulation in fentanyl vulnerability and suggest that enrichment-inspired, non-social interventions may offer a scalable strategy to reduce opioid use and relapse risk.

**Significance Statement:** This study demonstrates that long-term, object-based environmental enrichment in the absence of social peers is sufficient to reduce fentanyl intake and blunt stress-induced reinstatement in male and female rats. These findings identify a simple, scalable environmental intervention capable of dampening both drug use and vulnerability to relapse triggers. Given the prevalence of individuals exposed to periods of social isolation or with limited access to support, understanding how non-peer-based enrichment influences maladaptive opioid use and stress reactivity is of timely and translational importance. The ability of enrichment to reduce corticosterone, fentanyl consumption, and drug-cue reactivity highlights the importance of environmental context in shaping opioid misuse liability and implicates novel and practical avenues for relapse prevention

## Introduction

The current opioid epidemic remains a serious and escalating public health crisis. Since 2013, overdose has been the leading cause of injury death in the United States^1^. During the COVID-19 pandemic, opioid-related deaths rose sharply, increasing by 32% between 2019 and 2020^2^. Fentanyl accounted for the majority of these fatalities, with a 55% increase observed during the first year of the pandemic^3,4^. Beyond drug availability, pandemic-related factors - including social isolation, financial instability, and other stressors - disproportionately affected underserved communities^5–8^. These environmental stressors likely contributed to increased opioid use and relapse, emphasizing the need to understand how external stimuli shape opioid use, stress system function, and relapse vulnerability.

Stress is a well-established trigger for relapse. Early clinical studies demonstrated that relapse was positively associated with self-reported stressful life events following abstinence^9^. These observations were subsequently modeled in animals using the reinstatement paradigm, in which drug-seeking behavior is reinstated after abstinence or extinction by exposure to one of three relapse-inducing stimuli: a subthreshold drug dose, drug-associated cues, or stress^10–12^. A key shared feature among these stimuli is their capacity to activate the hypothalamic pituitary adrenal (HPA) axis, the central stress response system^13–16^. Accordingly, the stress-induced reinstatement procedure provides a valuable framework for examining neurobiological mechanisms underlying relapse. Traditional stressors in this model include footshock, forced swim, or tail pinch; however, pharmacological stressors such as kappa opioid receptor agonists, corticotropin-releasing factor, and the α2-adrenoceptor antagonist, yohimbine, are now more commonly used^16^. Yohimbine reliably induces HPA axis activation across species and readily instigates opioid-seeking behavior^11,15,17^. Therefore, yohimbine-mediated stress-induced reinstatement offers a translationally relevant approach for assessing the relationship between stress and relapse.

Environmental factors also have profound effects on drug craving and relapse. Environments associated with past drug use, or those lacking adequate health-related, financial, or educational resources, can provoke drug-seeking behavior in animal models and relapse in humans^14,18–24^. Conversely, environments that promote cognitive stimulation, social support, and/or physical activity can reduce drug craving and promote abstinence in both clinical and preclinical settings^25^. In rodent models, environmental enrichment (EE) is used to simulate positive and stimulating experiences by increasing the complexity of the living environment^26^. Effective and translational EE provides animals with opportunities for voluntary engagement in naturally rewarding, safe activities^26^, typically achieved by housing animals with multiple novel objects and at least one social peer, while control conditions typically include social-only housing (peers without objects) or isolated housing (neither peer nor objects)^27^. Numerous studies have reported the therapeutic effects of EE on behavioral measures related to drug abuse vulnerability^26,27^; however, these group-based enrichment conditions do not clarify whether beneficial effects of EE require both peers and objects or can be conferred by objects alone. This gap is particularly relevant given the increased prevalence of social isolation due to COVID-19 restrictions^28^ and the growing number of individuals working remotely or in hybrid environments^29^. Therefore, determining the therapeutic potential of enrichment in isolation is of timely importance.

Despite extensive work on EE, it has not been investigated whether object-based enrichment in isolation can reduce maladaptive fentanyl use or protect against stress-induced reinstatement of fentanyl-seeking behavior. Prior research shows that EE can lower baseline corticosterone levels – a physiological correlate of the canonical stress hormone cortisol in humans^30^ – and reduce reinstatement for both natural and drug rewards^26,31–34^. However, neither EE nor the stress-induced reinstatement model have been applied in the context of fentanyl. To address these gaps, the current study examined the efficacy of long-term, *non-peer, object-based* environmental enrichment in reducing fentanyl use and stress-induced relapse-like behavior in individually housed male and female rats. We further assessed whether EE altered baseline corticosterone concentrations and stress reactivity at key time points throughout the study. In the context of the current fentanyl crisis, these experiments aim to provide novel insight into how long-term enrichment in isolation impacts fentanyl use, reinstatement, and stress-reactivity.

## Materials and Methods

### Animals

Male and female wild-type Long-Evans Rats (n=37; 225 to 470g) were bred in house or purchased from Inotiv. Rats were given *libitum* access to water and were placed on a mild food restricted diet consisting of approximately 10% in body weight (g) of PicoLab® Rodent Diet 20 per day. Subjects were housed in temperature and humidity regulated conditions under a 12-hour light/dark cycle (lights on at 6:00 AM). Rats were group housed until they underwent intravenous catheterization and were individually housed thereafter. All animal procedures were performed in accordance with Washington University’s animal care committee’s regulations.

### Intravenous Catheterization surgery

To permit intravenous fentanyl self-administration, catheters were constructed in house with polyethylene tubing (∼13 cm, 0.02” ID, 0.037” OD) and were surgically implanted into the right jugular vein as previously described^35^. The tubing exited between the scapulae and was connected to a single-channel silicone vascular access harness (Instech Labs, VAHR1H/22). Enrofloxacin (8 mg/kg, s.c.) and carprofen (5 mg/kg, s.c.) and a chewable bacon-flavored rimadyl tablet (2 mg/tablet; Bio-Serv) were administered 0, 24, and 48 h after surgery. Catheter patency was maintained with daily flushing of 0.3 mL sterile saline containing gentamicin (1.33 mg/mL, i.v.) to mitigate infection. Patency was periodically assessed as needed with propofol (0.1 mL, 10 mg/mL, i.v.) and rats with a loss of patency at time point throughout self-administration were excluded from the study. Following catheterization, rats were randomly assigned to individually housed conditions with or without enrichment (see *Housing Conditions* below) for the duration of the study.

### Housing Conditions

Rats were assigned to either non-enriched (NE) or environmentally enriched (EE) single-housing conditions for the duration of the experiment (**Figure 1**; **Table 1**). All subjects were individually housed. In both NE and EE conditions, rats were housed in a standard durable, plexiglass cage with corn cob bedding, a stainless-steel chow and water rack, and a plastic, filtered cage top. NE conditions had no additional components. EE conditions contained five distinct enrichment items: (i) hand-selected foraging grass (Bio-Serv,) (ii) manzanita gnawing sticks (Bio-Serv), (iii) nylon bones (Bio-Serv), (iv) a primary novel enrichment item, (v) and a secondary enrichment item. Primary and secondary enrichment items were obtained from the [Fisher Scientific](https://www.fishersci.com/us/en/browse/90165027/animal-enrichment-devices?categoryKey=90165027&page=2) enrichment catalogue or fabricated using a 3D printer([Bambu/P1P](https://us.store.bambulab.com/collections/3d-printer?id=579551733527306247&skr=yes&gad_source=1&gclid=Cj0KCQjw782_BhDjARIsABTv_JAho9EGKdh417eBvnNnlT4VGRFlCbmMPjCRK9YXm5D5SwWRj14OTtMaAg8dEALw_wcB)) with lab grade polylactic acid (PLA) filament ([CARBON](https://www.amazon.com/Certified-Food-Grade-Printer-Filament/dp/B0BBBKJ6BY?ref_=ast_sto_dp&th=1)) (**Figure 1**). Primary items were categorized as one of the following functions: shelter (e.g., tunnel, hut), texture (e.g., dumbbell, Rubber chew), or interactive (e.g., grass woven swing, Stainless Steel Rattles). Secondary enrichment items consisted of a small unsophisticated object (e.g., kong toy, ping pong ball). Novel primary and secondary enrichment items were introduced during each cage change which occurred once per week independent of housing conditions (see **Table 1** below).

**Figure 1.**
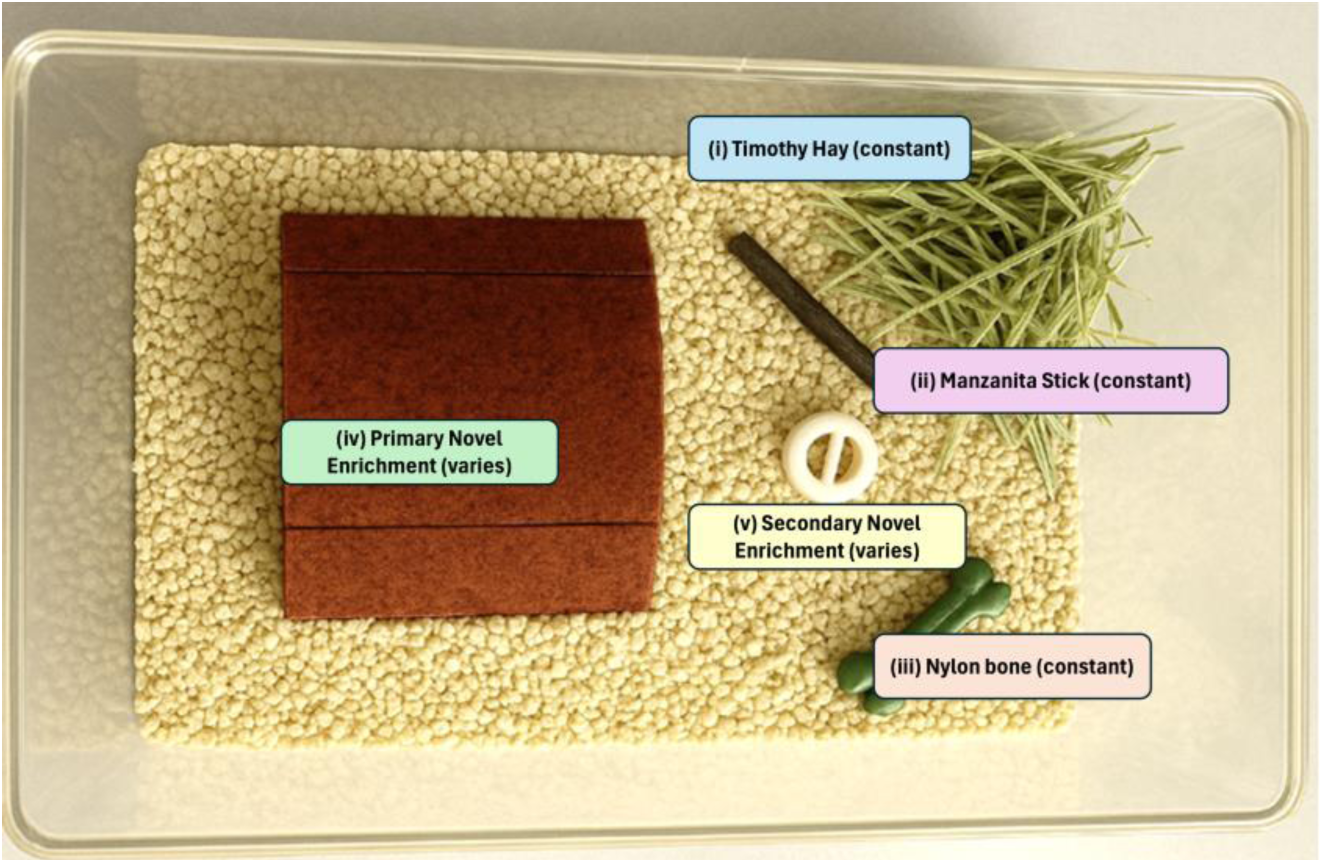
Example of environmentally enriched (EE) conditions consisting of five distinct stimuli (see **Table 1** for details).

**Table 1.** Environmental Enrichment.

| Item | Use | Type | Proposed Benefit | Source (Example) |
| --- | --- | --- | --- | --- |
| i | **Constant**; replaced each week | Timothy Hay | Encourages natural nesting and foraging behavior, aids in dental health | Bio-Serv #S1012 |
| ii | **Constant**; replaced each week | Manzanita Sticks | Promotes gnawing and aids in dental health, serves as perches or manipulanda | Bio-Serv #W0016 |
| iii | **Constant**; replaced each week | Nylon Bones | Promotes gnawing behavior and aids in dental health | Bio-Serv #K3580 |
| iv | **Primary Novel Enrichment**; type changes each week | Shelter Type | Enclosed space adds security, encourages nesting, reduces anxiety | Bio-Serv #K3365 (Rat-Hut); #K3325 (Rat-Tunnel); #K3245 (Rat Retreat) |
|  |  | Texture Type | Promotes exploratory behavior; encourages forelimb use, reduces boredom | Bio-Serv #K322 (Dumbbells); #K3289 (DNA Flexor); #K3290 (Football); #K341 (Jump'n Jacks) |
|  |  | Interactive Type | Encourages movement; provides mental stimulation; engages curiosity through object novelty and portability | Fisher Scientific #14-727-066 (rattles); #14-727-032 (ball and chain); Rat foraging Toy 3d Print File |
| v | **Secondary Novel**; similar type changed each week | Non-specific unsophisticated | introduces novelty; serves as additional manipulanda with portability | Airless Golfball 3D Print File<br>Rattling Ball 3D print file |

### Drugs

Fentanyl stock solution (50 μg/mL) was diluted with a sterile saline solution to obtain the appropriate dose based on body weight (5 or 2 ug/kg/100 μL infusion). Yohimbine ([ThermoFisher](https://www.thermofisher.com/order/catalog/product/141030050)) was dissolved in a 1:1 solution of sterile deionized (DI) water and 0.9% saline using sonication, yielding a final concentration of 2.5 mg/mL.

### Fentanyl Self-Administration, Extinction, and Reinstatement

The fentanyl self-administration, extinction, and reinstatement procedure was conducted as previously described^35^. All procedures were performed in a standard operant chamber (Med Associates, RRID: SCR_021938) equipped with two retractable levers, a cue light above each lever, a stainless-steel grid flooring, and a house light. Rats were naive to the apparatus and instrumental training prior to the start of self-administration. For fentanyl self-administration (SA) sessions, rats were placed in operant chambers and tethered to the infusion line. Rats underwent fifteen 2-hour SA training sessions under a fixed-ratio 1 (FR1) reinforcement schedule. Responses on a designated active lever resulted in illumination of the light cue above the active lever (5 s) and the programmed delivery of a single fentanyl infusion (100 μL, 5.24 s) followed by a 20-sec time-out period during which additional active lever responses did not result in added reinforcement. Responses on the inactive lever were recorded but had no program consequences. Rats were allowed to self-administer a moderate dose of fentanyl dissolved in sterile saline during the first 5 sessions of Acquisition (5 μg/kg/infusion, i.v.) and had access to a lower dose the remaining 10 sessions during Maintenance (2 μg/kg/infusion, i.v.) as previously described^35^.

Upon completing 15 SA sessions, rats underwent extinction training which took place over the course of seven 1-hour sessions in the same operant chamber as SA except that rats were not tethered to the infusion line and no associated cues or rewards were administered following active and inactive lever responses. On the last day of extinction training, rats were injected with saline (1 mL/kg, i.p.) 20 minutes before the start of session to habituate them to the injection procedure performed during the subsequent reinstatement session (see behavioral timeline in **Figure 2A**).

**Figure 2.**
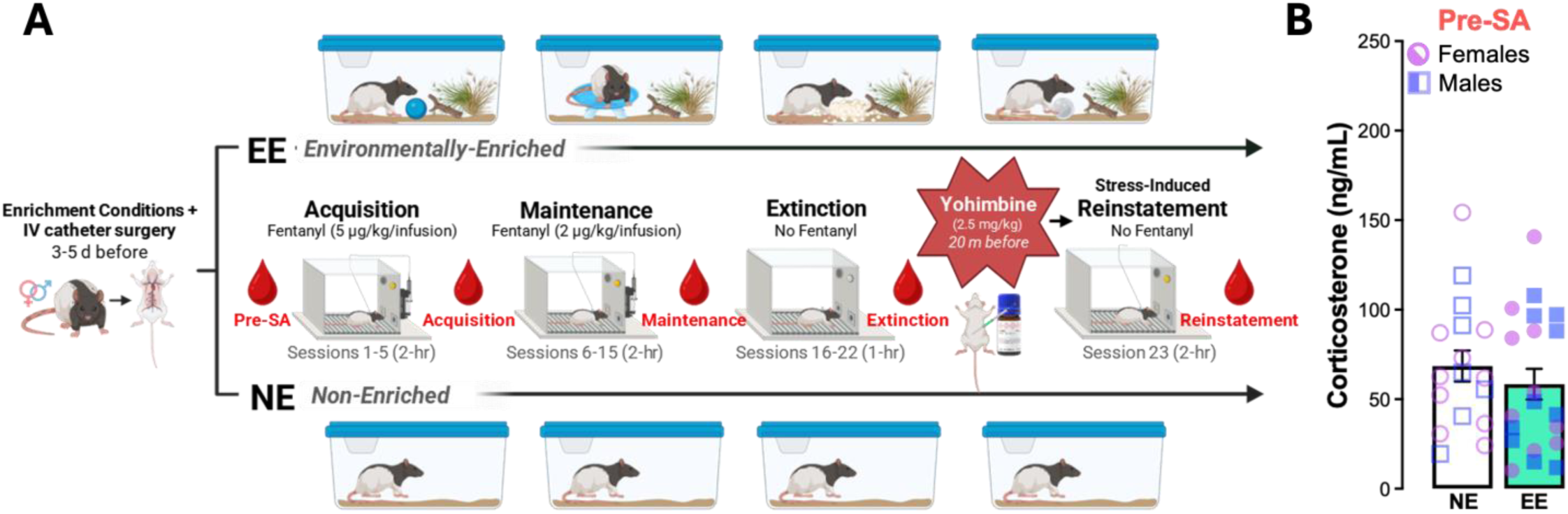
Experimental timeline and baseline corticosterone concentrations. **(A)** Male and female Long-Evans rats received intravenous (IV) jugular catheters 3-5 d prior to the first day of fentanyl self-administration. After surgery, rats were randomly assigned to be individually housed with environmental enrichment (EE) or no enrichment (NE) which were maintained over the course of the experiments. Acquisition of fentanyl self-administration occurred over 5 2-hr sessions where responses on an active lever triggered a light cue illumination and delivery of a fentanyl infusion (5 µg/kg, IV). Subsequent maintenance sessions (6-15) occurred in the same manner except with a lower fentanyl dose of 2 µg/kg, IV. Upon completing the 15 sessions of fentanyl self-administration, rats underwent 7 1-hr sessions of extinction training where responses on the active and inactive lever were recorded but had no programmed consequences. The next day, rats received yohimbine (2.5 mg/kg, i.p.) 20 minutes before the 2-hr reinstatement test during which conditions were identical to those during self-administration except fentanyl reinforcement was withheld. Tail vein blood samples were obtained at multiple time points throughout the study (red): one day before starting fentanyl self-administration (Pre-SA), after sessions 3 or 4 (acquisition), after one of sessions 8-12 (maintenance), after session 21 or 22 (extinction), and trunk blood was obtained immediately after the final test (reinstatement). **(B)** Prior to starting self-administration (Pre-SA), EE did not alter baseline corticosterone levels (unpaired t-test). N=18-19/group; Females depicted as circles, males depicted as squares.

For the final test of stress-induced reinstatement, rats received the pharmacological stressor, yohimbine (2.5, mg/kg, i.p.) 20 minutes before the start of the test. Reinstatement took place in the same operant apparatus as SA and extinction. During the two-hour test, conditions were identical to those present during SA training except that fentanyl reinforcement was withheld. Responses on the active lever were used as an indicator of drug-seeking behavior. Immediately following completion of the 2-hour test, rats were deeply anesthetized, decapitated, and trunk blood was obtained in leu of tail blood (see below) to assess corticosterone concentrations during reinstatement.

### Tail Blood Collection

Blood samples were obtained between 11:00 AM and 4:00 PM upon completing behavioral sessions at 5 time points throughout the experiment (see **Figure 2A**): (1) one day prior to the first fentanyl SA session to establish baseline corticosterone levels (Pre-SA), (2) after one of the first 5 fentanyl SA sessions (Acquisition), (3) after one of SA sessions 8-12 (Maintenance), (4) once after extinction session 6 or 7 (Extinction) and (5) after administration of yohimbine and the reinstatement test (Reinstatement). All blood samples (except Reinstatement) were obtained vial tail nick of the lateral tail vein as previously reported^14^. Briefly, rats were swaddled, and the distal end of the rat’s tail was secured while a sterile razor was used to nick <1 cm parallel to the lateral tail vein. The tail was milked from the proximal to the distal end using the thumb and index finger and blood samples were collected in serum separating tubes. Samples coagulated at room temperature for one hour before they were centrifuged for 15 minutes at 4 degrees C° at 13,000 RPM. The separated serum was decanted and aliquots of 25 μL-50 μL were prepared and stored at -80 C° until assayed.

### Corticosterone Analyses

Serum corticosterone concentrations were determined using a competitive enzyme immunoassay ELISA kit (Bio-Techne; sensitivity=0.047 ng/mL, range=0.103-25 ng/mL) according to the assay instructions. Briefly, samples were thawed on ice, pretreated, and diluted 8x with calibrator diluent. Duplicates were run from each sample and standard on multiple plates. The pre-treated 96-well plate was incubated with a polyclonal antibody specific for corticosterone and a substrate solution was added to the wells to determine the bound enzyme activity determined by the absorbance. Plates were read at 450 nm within 15 minutes of development. Duplicate readings for each standard and sample were averaged, and the optical density (OD) of non-specific binding (NSB) was subtracted. A standard curve for each plate was generated using a four parameter logistic (4-PL) curve (R^2^=0.979-0.991). Then corticosterone concentrations were determined based on the interpolated value from the standard curve and multiplied by the dilution factor. Values outside the range of detectability were excluded from the analysis.

### Statistical Analysis

All statistical analyses were conducted using GraphPad Prism. Data were assessed for normality and sphericity; Greenhouse–Geisser corrections were applied where necessary. Behavioral data from self-administration, extinction, and reinstatement phases were analyzed using repeated-measures 3-way ANOVAs with session as the within-subjects factor and housing condition (environmental enrichment [EE] vs. non-enrichment [NE]) and sex (male vs. female) as between-subjects factors. There were no effects of sex in these analyses (see **Table 2**), thus statistical comparisons were carried out with repeated-measures 2-way ANOVAs. Where appropriate, Tukey’s and Sidak’s multiple comparisons tests were used post-hoc. Based on the lack of sex effect in behavioral findings, corticosterone concentrations were assessed between enrichment groups using unpaired t-tests and simple linear regressions for each time point and repeated measures 2-way ANOVAs for comparisons across different time points. Comprehensive reporting of all statistical analyses performed are in **Table 2**. Statistical significance was set at α = 0.05 for all tests. Data are presented as mean ± SEM.

**Table 2.**
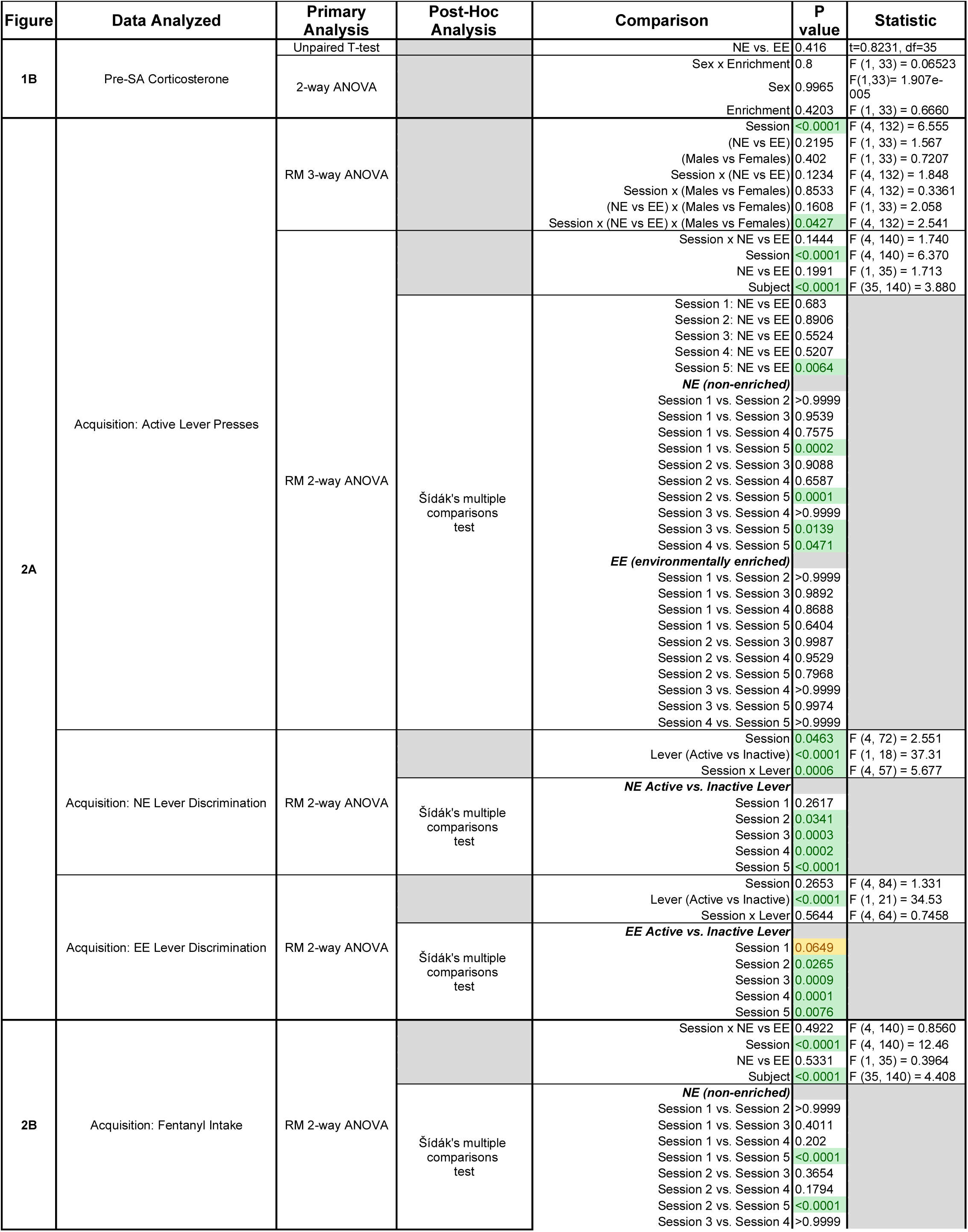
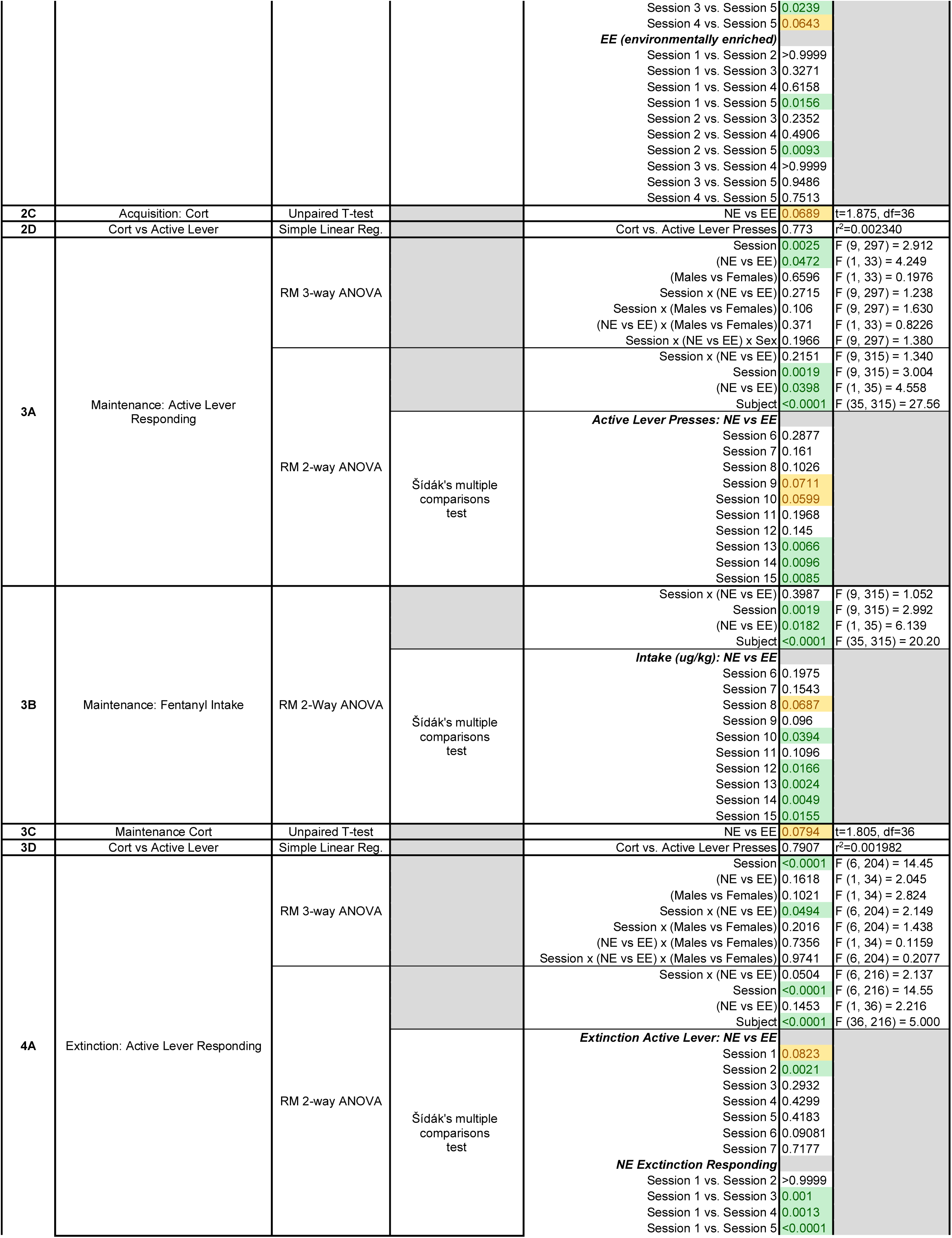
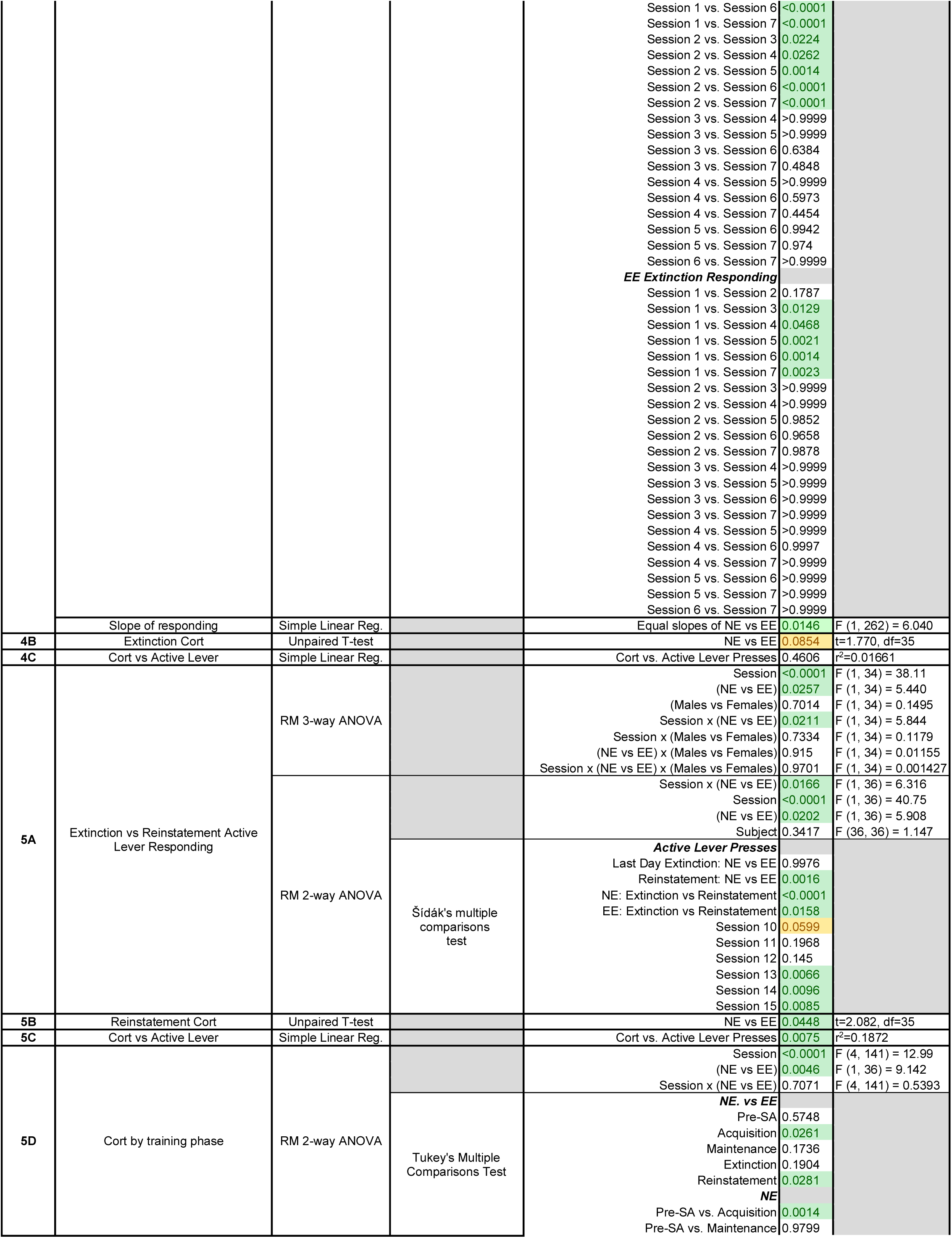
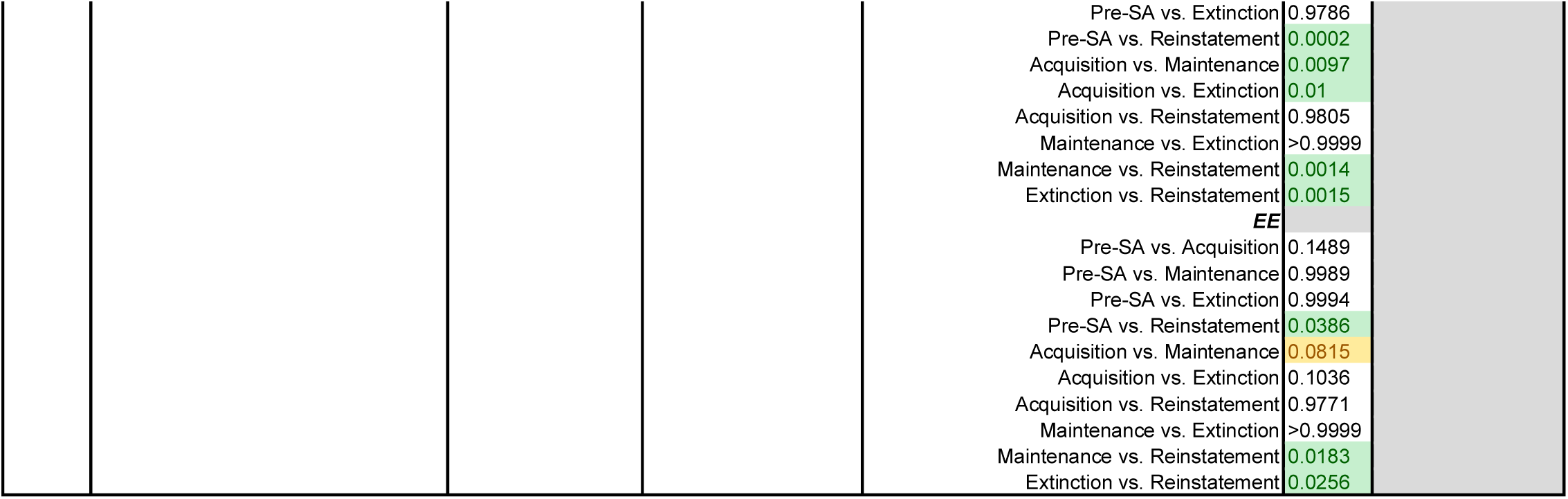
Detailed statistical table.

## Results

### Prior to fentanyl self-administration, environmental enrichment in isolation does not alter baseline corticosterone concentrations

Following IV catheterization surgery and postoperative care, subjects were acclimated to their non-enriched (NE) or environmentally enriched (EE) isolated housing conditions (**Figure 1**) for at least three days before tail blood samples were obtained to assess baseline differences in corticosterone concentrations. Prior to starting fentanyl self-administration (Pre-SA), there were no differences in corticosterone concentrations between EE and NE groups (**Figure 2B**) indicating that short-term EE (3-5 days) in the absence of fentanyl exposure does not impact baseline HPA axis activation.

### Environmental enrichment alters the trajectory of acquisition of fentanyl self-administration

Rats received no prior training or exposure to the operant chambers prior to starting fentanyl self-administration. During acquisition, rats underwent five 2-hour sessions of fentanyl self-administration (5 µg/kg/infusion; **Figure 2A**). Interestingly, housing enrichment conditions did not impact the ability of rats to discriminate between the active and inactive lever and both groups readily acquired fentanyl self-administration regardless of sex. However, EE influenced the rate of responding during this initial stage of training because NE rats responded more than EE rats on the last day of acquisition (session 5) and more than previous days of training (**Figure 3A**). By contrast, EE rats maintained stable responding over the five days of acquisition. Nonetheless, fentanyl intake across the five sessions remained similar between groups (**Figure 3B)**, suggesting that NE rats responded more during time-out periods, possibly indicating higher impulsivity. Although not significant, we also observed a trending decrease in EE corticosterone levels compared to NE (**Figure 3C**), but these were unrelated to lever responding during acquisition (**Figure 3D**). Given that differences between EE and NE behavior was not evident until five self-administration sessions, these findings suggest that environmental enrichment does not alter the ability to acquire drug-cue associations but may rather reduce the salience of drug cues or drug reinforcement.

**Figure 3.**
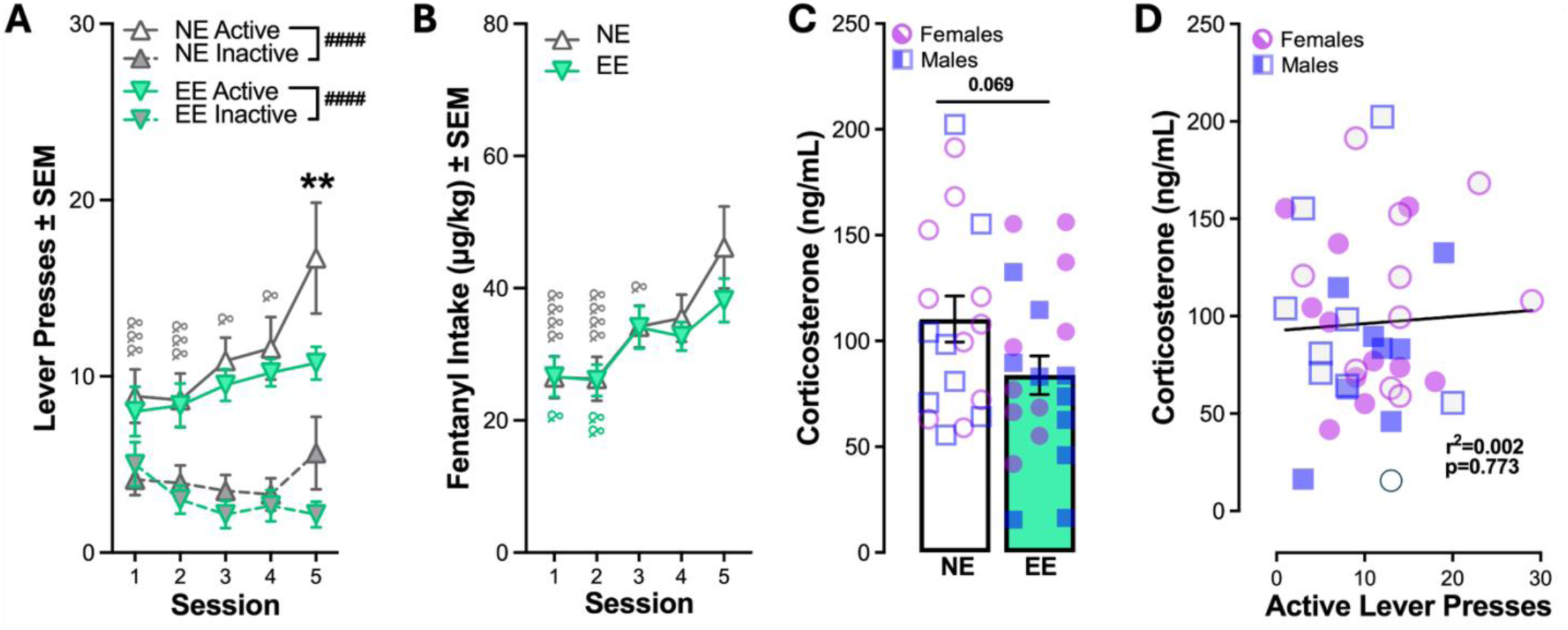
Acquisition of fentanyl self-administration is altered by environmental enrichment. **(A)** Both environmentally enriched (EE, n=18) and non-enriched (NE=19) groups acquire fentanyl self-administration and discriminate between the active and inactive lever. The trajectory of responding varied by housing condition in that NE rats responded more on the active lever the last day of acquisition than EE rats (**) and more than any previous day of training (&). **(B)** Fentanyl intake was similar between NE and EE groups, increasing over time. **(C)** A trending decrease in corticosterone concentrations was observed in EE rats compared to NE rats during acquisition but **(D)** corticosterone levels were not related to responding during training. RM 2-way ANOVAs and Sidak’s multiple comparisons tests (A-B), unpaired t-test (C), and simple linear regression (D). N=18-19/group; Females depicted as circles, males depicted as squares, and both depicted as triangles. Symbols indicate **#**main effects of lever, *between-group differences, and **&**within-group differences relative to session 5. (*p<0.05, **p<0.01, ***p<0.001, ****p<0.0001). Full statistical reporting available in **Table 2**.

### Environmental enrichment attenuates the maintenance of fentanyl self-administration responding and intake

Maintenance of fentanyl self-administration continued in ten subsequent 2-h sessions except that the dose of fentanyl was lower (2 µg/kg/infusion). Independent of sex, both groups continued to preferentially respond on the active lever, NE rats made more responses for fentanyl than EE rats, particularly on the last three days of self-administration (**Figure 4A**). Similarly, fentanyl intake was, overall, higher in NE rats relative to EE (**Figure 4B**), demonstrating that EE suppressed fentanyl use. Similar to the observed effects during acquisition, we found a trending decrease in corticosterone levels in EE rats compared to NE (**Figure 4C**), but these were unrelated to responding during the maintenance session for which samples were obtained (**Figure 4D**).

**Figure 4.**
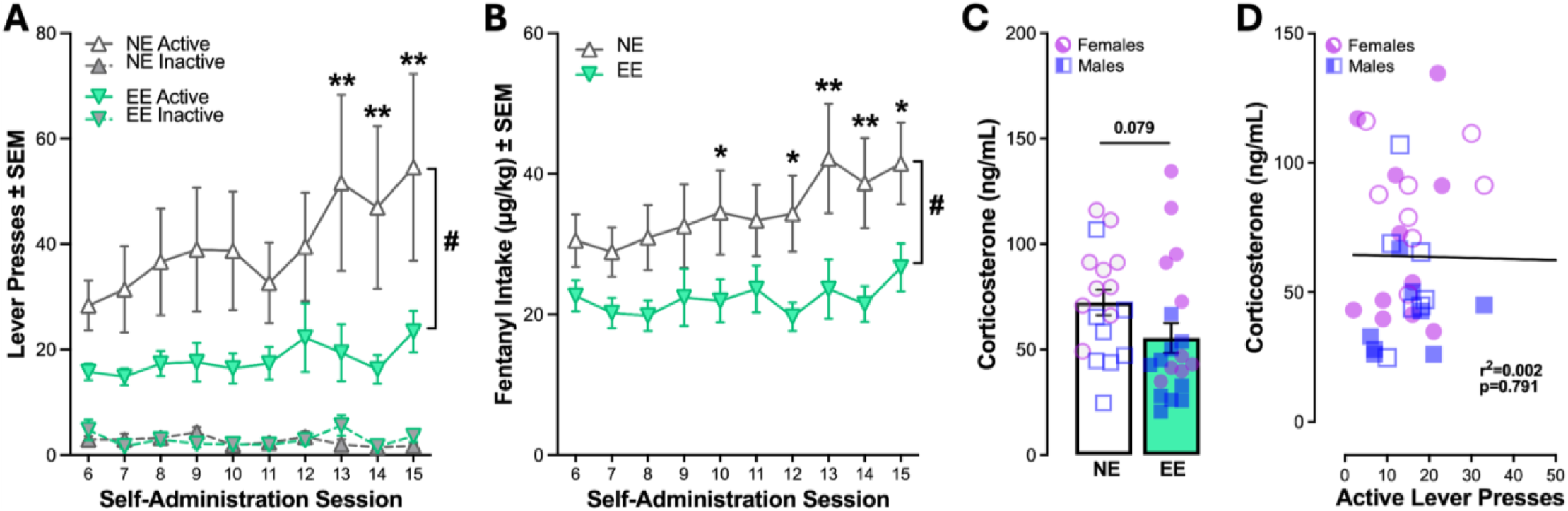
Maintenance of fentanyl self-administration is attenuated by environmental enrichment. **(A)** Active and inactive lever pressing during maintenance of fentanyl self-administration (sessions 6-15). EE attenuates active lever responding for fentanyl and **(B)** attenuates fentanyl intake. **(C)** A trending decrease in corticosterone concentrations was observed in EE rats compared to NE rats during maintenance but **(D)** corticosterone levels were not related to responding during training. RM 2-way ANOVAs and Sidak’s multiple comparisons tests (A-B), unpaired t-test (C), and simple linear regression (D). N=18 -19/group; Females depicted as circles, males depicted as squares, and both depicted as triangles. Symbols indicate **#**main effects of housing and *between-group differences. (*p<0.05, **p<0.01, ***p<0.001, ****p<0.0001). Full statistical reporting available in **Table 2**.

### Environmental enrichment facilitates extinction of drug-seeking during extinction training

After completing acquisition and maintenance of fentanyl self-administration, rats underwent seven 1-hr sessions of extinction training where fentanyl and cue reinforcement was withheld. Independent of housing conditions or sex, rats reduced their responding on the active lever over the seven days of training (**Figure 5A**). However, EE impacted the rate of extinction. NE rats continued to exhibit high rates of responding during the first two sessions compared to subsequent extinction sessions while EE rats only showed higher drug-seeking on day 1 (see **Table 2**). Analysis of active lever responding across the seven days revealed significant differences in the rate of extinction between NE and EE rats, indicating that NE rats show more drug-seeking behavior at earlier timepoints. This suggests that EE reduces the persistence of drug-seeking in the absence of drug-related cues. Nevertheless, these behavioral differences did not translate to differences in corticosterone concentrations (**Figure 5B-C**).

**Figure 5.**
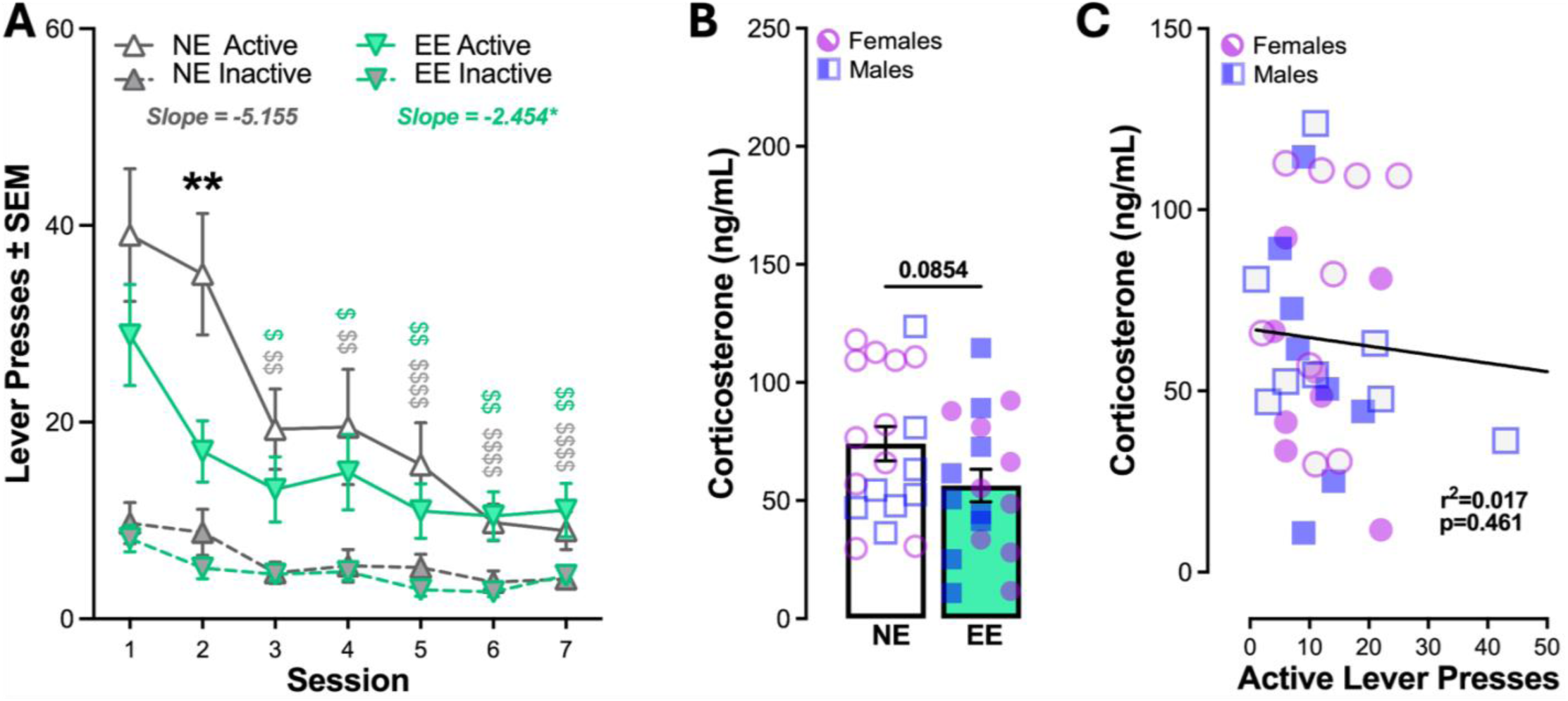
Extinction training is impacted by environmental enrichment. **(A)** Active and inactive lever pressing during extinction (when fentanyl and cues are absent). NE rats respond more than EE rats on day 2 of extinction (**), but both groups reduce responding over time ($relative to day 1). Simple linear regression of active lever responding during extinction revealed significant differences in the slope between NE and EE. **(B)** A trending decrease in corticosterone concentrations was observed in EE rats compared to NE rats during extinction but **(C)** corticosterone levels were not related to responding during training. RM 2-way ANOVAs and Sidak’s multiple comparisons tests (A), unpaired t-test (B), and simple linear regression (C). N=18-19/group; Females depicted as circles, males depicted as squares, and both depicted as triangles. Symbols indicate **#**main effects of housing and *between-group differences. (*p<0.05, **p<0.01, ***p<0.001, ****p<0.0001). Full statistical reporting available in **Table 2**.

### Environmental enrichment attenuates stress-induced reinstatement of fentanyl-seeking behavior and associated stress-reactivity

The next day after the last extinction session, rats underwent a test of stress-induced reinstatement precipitated by Yohimbine (2.5 mg/kg, i.p.) administered 20 min before the session. During the test, rats were tethered to the infusion line but fentanyl reinforcement was withheld. Active lever responses during the test were used as a proxy of drug-seeking behavior. Independent of housing conditions, rats reinstated fentanyl-seeking behavior as indicated by a significant increase in active lever responses relative to the previous day in extinction (**Figure 6A**). However, the magnitude of reinstatement was greater in NE rats relative to EE rats, demonstrating that EE attenuates stress-induced drug-seeking behavior. This effect was likely attributed to the corresponding reduction in stress reactivity as indicated by lower corticosterone concentrations observed in EE rats compared to NE (**Figure 6B**). Moreover, corticosterone was positively associated with active lever pressing during the session (**Figure 6C**), suggesting the effects on behavior were related to stress levels instigated by yohimbine. Assessment of group corticosterone levels throughout the experimental timeline revealed that EE resulted in, overall, lower corticosterone levels (**Figure 6D**). Interestingly, NE and EE differences were most evident during acquisition and reinstatement, but only reinstatement was significantly higher than pre-SA levels in both groups. As such, higher corticosterone levels in NE groups during acquisition and maintenance may reflect EE-dependent alterations in stress reactivity, potentially provoked under conditions of uncertainty (i.e., novel conditions). Together, these findings demonstrate the efficacy of object-based enrichment in isolation in reducing fentanyl use and stress-induced drug craving.

**Figure 6.**
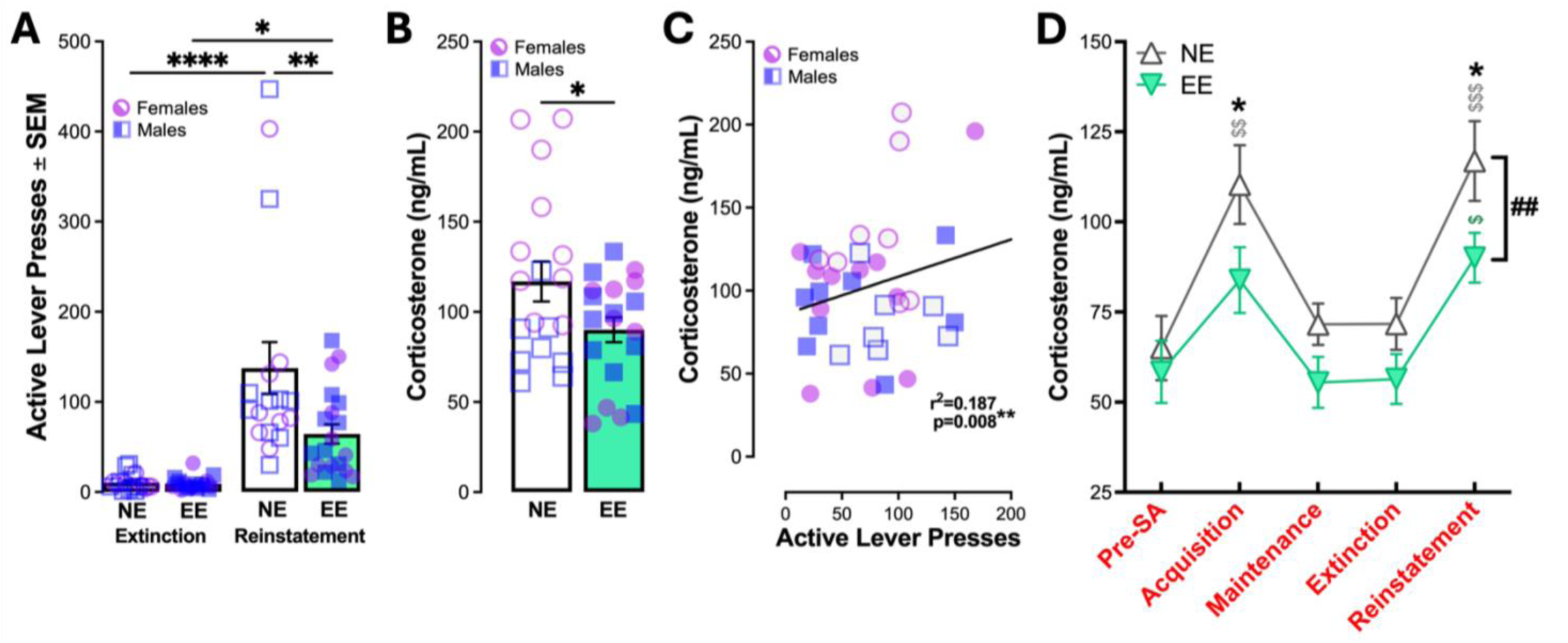
Environmental enrichment attenuates stress-induced reinstatement and associated stress reactivity. **(A)** Yohimbine (2.5 mg/kg) was administered 20 min before the 2-hr reinstatement test. Both groups reinstated based on an increase in active lever pressing during reinstatement relative to extinction. Reinstatement was lower in rats with EE compared to NE. lever pressing on the last day of extinction and during reinstatement. (RM 2-way ANOVA, *Sidak’s multiple comparisons test). **(B)** Corticosterone concentrations after reinstatement were lower in rats with EE relative to NE (*Unpaired t-test) and **(C)** Corticosterone levels are positively associated with active lever responding during the reinstatement test (*Simple linear regression). **(D)** EE produces an overall reduction in corticosterone levels which were significantly less than NE during acquisition and maintenance. NE levels were higher than baseline (Pre-SA) during acquisition and reinstatement, but EE levels were only higher during reinstatement (RM 2-way ANOVA, ^#^main effect of housing; Tukey’s multiple comparisons test, *between group difference, ^$^within-group difference relative to Pre-SA). N=18-19/group; Females depicted as circles, males depicted as squares, and both depicted as triangles. (*p<0.05, **p<0.01, ***p<0.001, ****p<0.0001). Full statistical reporting available in **Table 2**.

## Discussion

The current study demonstrates that long-term, non-social, object-based EE attenuates fentanyl self-administration, reduces stress reactivity, and limits stress-induced drug-seeking behavior in both male and female rats. These findings indicate that enrichment delivered in the absence of social peers is sufficient to blunt maladaptive opioid use and stress-induced relapse-like behavior. These add to a substantial body of evidence indicating that environmental enrichment can mitigate addiction-like behaviors across drug classes^31,36,37^, and importantly, are the first to show such effects in the context of fentanyl – currently the most lethal contributor to overdose mortality.

### Environmental enrichment reduces fentanyl-taking and drug-seeking

Consistent with prior work showing that EE decreases self-administration of psychostimulants, opioids, and alcohol^38–40^, we found that EE suppressed fentanyl self-administration during both acquisition and maintenance. NE rats exhibited increased active lever responding toward the end of acquisition (**Figure 3A**) that continued into maintenance (**Figure 4**), while EE rats maintained a stable pattern of drug use and consumed less fentanyl during training. Importantly, housing conditions did not influence the ability of rats to acquire fentanyl self-administration (**Figure 3**) similar to previous findings with amphetamine, cocaine, and nicotine^41–43^. The difference in the trajectory suggests that EE may reduce the reinforcing efficacy or incentive salience of fentanyl or fentanyl-associated cues^44^. Previous studies have proposed several non-mutually exclusive mechanisms underlying this effect, including: increased engagement with alternative non-drug rewards^45–47^, altered mesolimbic dopamine function^48–50^, and enhanced synaptic plasticity within prefrontal corticostriatal circuits^51,52^.

Our design implements enrichment as object-based novelty and interaction rather than social complexity, a distinction that has been rarely addressed in past research. The fact that enrichment without social peer interaction was sufficient to suppress fentanyl seeking implies that physical and cognitive stimulation – rather than social buffering alone – can shift the valuation of drug reinforcers. Several lines of evidence confer the detrimental effects of social isolation^53^ and the benefits of environmental enrichment on maladaptive drug use^46^. However, the latter is largely based on effects derived from both peer- and object-based enrichment and therefore fail to discern the efficacy of object-based enrichment independently. Notably, one study examined long- and short-access cocaine self-administration in rats housed in (a) social conditions with object enrichment, (b) social conditions without objects, (c) isolated conditions with object enrichment, and (d) isolated conditions^43^. Although not directly compared, the authors showed that rats in isolated conditions with object enrichment self-administered cocaine at levels similar to those with social and object enrichment^43^. Similarly, a separate study found that pair-housed rats with EE exhibited less cocaine-seeking behavior in extinction that pair-housed rats without EE^54^. Based on these in the context of the current findings, the therapeutic potential of enrichment on addictive behaviors may rely more on object-based novelty and interaction. This notion is particularly relevant for individuals facing isolation due to pandemic-related disruptions or remote work environments.

### Enrichment reduces persistence of drug-seeking during extinction

Although both NE and EE groups extinguished responding across sessions, NE rats exhibited greater persistence in drug-seeking during early extinction sessions (**Figure 5**). This aligns with the idea that enrichment enhances cognitive flexibility and accelerates extinction learning^32,54,55^. Reduced perseveration suggests that EE rats adapt more readily to the absence of previously learned reinforcement contingencies. Mechanistically, enrichment has been shown to enhance prefrontal cortical function, increase dendritic branching, and upregulate plasticity markers, like BDNF^52,56,57^. These neuroadaptations may facilitate inhibitory control over drug-seeking responses when drug-related contingencies change.

### Enrichment attenuates stress-induced reinstatement

Strikingly, EE blunted yohimbine-induced reinstatement of fentanyl seeking (**Figure 6**). Stress is a primary driver of relapse in humans^58^, and yohimbine-induced reinstatement robustly models stress-precipitated craving via engagement of the HPA axis^59^. Our results showing that NE rats exhibit stronger reinstatement than EE rats, indicate that enrichment modulates vulnerability to stress-triggered relapse-like behavior. Corticosterone analyses further support a stress-buffering role for enrichment. EE rats had lower corticosterone concentrations after reinstatement, and across the entire experiment (**Figure 6B-D**), suggesting that EE reduced HPA axis responsivity. This aligns with past findings that long-term EE lowers basal glucocorticoid levels and dampens physiological stress reactivity^30^. Importantly, corticosterone was positively correlated with reinstatement responding, suggesting that enriched environments reduce relapse-like behavior, in part, by constraining stress system activation. This interpretation aligns with studies showing that EE reduces anxiety-like behavior, modifies CRF and glucocorticoid receptor expression, and enhances stress coping strategies^60,61^.

### Enrichment effects in social isolation: implications and mechanisms

By using individually housed rats in both NE and EE conditions, the current study isolates object-based enrichment effects from potential confounds of social interaction. Surprisingly, few studies have systematically separated these components. Our data show that the therapeutic benefits of enrichment persist even in the absence of social peers, suggesting that cognitive, sensory, and motor engagement alone deliver sufficient buffering against maladaptive opioid use and stress-induced drug craving. This distinction is critical given the increasing social isolation across populations and the need for accessible, non-social forms of environmental novelty and stimulation. The protective effects of object-only enrichment may arise from engagement with naturalistic behaviors (gnawing, exploring, manipulating), which can reduce boredom, anxiety, and passivity – factors associated with drug abuse vulnerability^62^. Moreover, we observed no sex differences in fentanyl self-administration, extinction, or reinstatement. While some studies report female-specific vulnerabilities to opioid and psychostimulant reinforcement^63^, others find minimal sex differences when access, dosing, and stress variables and tightly controlled^64,65^. The absence of sex effects in the current study suggests that the protective impact of object-based EE generalizes across sexes in this paradigm.

### Limitations and future directions

Several limitations warrant consideration. First, although EE reduced reinstatement triggered by yohimbine, it remains unknown whether similar protective effects would extend to cue-induced, drug-primed, or alternative stressor-induced reinstatement. Given that EE may influence cue salience and reward valuation, it will be important to test EE in isolation with other relapse modalities. Second, while corticosterone provides a meaningful measure of HPA axis activity, other neurobiological readouts of stress system function like CRF expression or noradrenergic activity may clarify more detailed underlying mechanisms. Finally, although singly housed enrichment allowed isolation of object-based effects, social enrichment remains a powerful determinant of addiction vulnerability, and future work should directly compare object-only, social-only, and combined enrichment approaches.

### Conclusions

Our findings show that long-term, object-based environmental enrichment is sufficient to blunt fentanyl self-administration, reduce the persistence of drug-seeking, and protect against stress-induced reinstatement and stress reactivity. These results highlight the power of environmental context – and specifically, access to stimulating, engaging, and rewarding non-social activities – to alter opioid abuse vulnerability. In the context of the ongoing fentanyl crisis, these data underscore the potential translational utility of enrichment-inspired interventions that are feasible in socially isolated or resource-limited settings. Such interventions may provide a low-risk and scalable means to reduce opioid use and relapse risk, complementing existing pharmacological and behavioral therapies.

## Acknowledgements

The authors thank Justin Meyer for breeding colonies and general managerial support. This work was supported by NIH DA063220 (JAM), DA054900 (JAM), DA058613 (JAM).

